# Accelerated discovery of thermostable vaccines using data-efficient AI

**DOI:** 10.64898/2026.09.17.752370

**Authors:** Jinbi Tian, Khanh T.M. Tran, Brett H. Pogostin, Olivia Sheridan, Sevinj Mursalova, Amy H. Lee, Shuai Liu, Jaya Hamkins, Daniel Antov, Alana L. Power, Zane S. Dash, Dongsoo Yun, Mina Konaković Luković, Robert S. Langer, Ana Jaklenec

**Author notes:** Corresponding author. (R.L.), (A.J.). These authors contributed equally to this work (Jinbi Tian, Khanh T.M. Tran).

## Abstract

The inherent instability of mRNA-lipid nanoparticles (LNPs) necessitates ultra-cold storage, creating significant barriers for global distribution and limiting their broader application in advanced delivery systems. Solid-state, water-free formulations offer a promising solution by enhancing thermostability and enabling integration into emerging delivery modalities such as microneedle (MN) patches. Prior efforts to stabilize mRNA-LNPs have been constrained by narrow formulation scope and low-throughput screening methods. Here, we introduce AGENT (Algorithm-Guided Experimental design for lipid Nanoparticle Thermostabilization), an AI-driven framework that couples high-throughput experimentation with Bayesian optimization to rapidly identify thermostable mRNA-LNP formulations. Manual exploration of the formulation space required months of screening and yielded suboptimal candidates. In contrast, AGENT extracted maximal information from sparse experimental datasets, enabling efficient formulation optimization in only six iterations completed within one month. Using AGENT, we stabilized mRNA vaccines with diverse LNPs, including those in clinical use, into solid state formulations that retained 100% bioactivity after storage at 37°C for over two months. The thermostable vaccines induced antigen-specific IgG and germinal center B cell responses that were non-inferior to those elicited by freshly prepared soluble vaccines. The solid-state formulations were further incorporated into dissolvable MN patches and administered to rodents and nonhuman primates, yielding comparable neutralizing antibody titers compared to conventional intramuscular delivery of fresh vaccines. To our knowledge, this study presents the first demonstration of AI-driven design of thermostable RNA vaccines, offering a scalable, cold-chain-free solution for global immunization. By addressing both stability and delivery challenges, AGENT provides a potentially transformative platform for developing accessible next-generation therapeutics.

## Introduction

Lipid nanoparticles (LNP) have advanced RNA technology to the forefront of prophylactics and therapeutics development for infectious diseases, cancers, and autoimmune conditions^1,2^. While mRNA-LNPs offer advantages such as safety, scalability, and design flexibility, they are inherently unstable^3–5^. Consequently, stringent storage conditions and the incorporation of appropriate excipients are essential for maintaining the product’s integrity during the manufacturing process and throughout its life cycle. Yet, their shelf life remains short. For examples, the recommended storage for a commercial COVID-19 vaccine is 60 days at 2-8°C and 10 months at-50°C to-15°C^6^. Although the cold chain ensures vaccines potency, the ultra-cold temperature requirement is a significant barrier to global vaccine administration and distribution, especially in limited-resource settings^7,8^. Primary wastage, estimated to be 30% to 40% in the total cost of immunization, comes from improper vaccine storage and expiration^9^. Additionally, the instability of mRNA-LNPs limit their applications in advanced controlled-delivery systems that could improve access, distribution and durability. Therefore, there remains a critical need to improve mRNA-LNP stability and eliminate ultra-cold storages.

An effective approach to enhance the thermostability of mRNA-LNPs is to develop solid-state, water-free formulations. By minimizing physical stresses associated with aqueous environments, such as changes in pH, ionic strength, agitation, interface exposure, and hydrolytic degradation^10–12^, solid-state formulations offer longer shelf-life and more flexibility in transportation and storage compared to liquid formulations^13–15^. Moreover, solid-state mRNA-LNPs can be integrated into advanced drug delivery devices, such as microneedle (MN) patches, which offer needle-free, pain-free delivery, simplify logistics by eliminating sharp waste and the need for trained personnel^16–18^. As such, MN hold particular promise for improving compliance and expanding vaccine access in low-resource settings. To fully realize the potential of MN-administered vaccines, mRNA-LNPs must be stabilized in solid forms. While excipient-based stabilization has shown promise^14,15,19–22^, past efforts were confined to a limited range of LNP chemistries and small-scale manual screenings, leaving much of the design space unexplored. Furthermore, recent attempts to develop MN patches for mRNA delivery using commercial LNP compositions have highlighted the difficulty of stabilizing these formulations^23,24^, revealing a significant gap in existing approaches and the need for a more comprehensive approach.

The rapid rise of artificial intelligence (AI) has profoundly reshaped the discovery and development pipeline across the life sciences. In drug formulation, a central challenge is the slow and resource-intensive screening of candidates and process conditions^25,26^. AI offers a paradigm shift, replacing arduous trial-and-error screening with efficient predictive, data-driven exploration. However, most AI and deep learning approaches require large, curated datasets^27^, a barrier in emerging fields such as LNP solid-state formulation, where prior knowledge is scarce and experimental throughput is constrained. Motivated by these challenges, we developed AGENT (Algorithm-Guided Experimental design for lipid Nanoparticle Thermostabilization), a framework that couples high-throughput screening with Bayesian optimization. AGENT guides the data collection in a sample-efficient way, leveraging targeted wet-lab data to iteratively predict and refine formulations, compressing the development cycle from months or years to days.

In this study, we examined AGENT’s versatility in stabilizing various LNP compositions, including those already used in commercial vaccines. The stabilized formulations remained bioactive at 37°C for at least two months, which eliminated the need for cold-chain storage. Additionally, the AI-discovered thermostable formulations were adapted to MN patch delivery and successfully employed in rodent and non-human primate models, validating our AI approach for thermostable solid-state mRNA-LNPs. To our knowledge, this is the first use of AI to accelerate the design of thermostable RNA-based vaccines. By integrating empirical testing with algorithmic insight, AGENT provides a powerful platform for creating globally accessible vaccines. The combination of thermostable solid-state design and MN patch delivery addresses both stability and administration barriers, paving the way for equitable distribution of next-generation RNA-based medicines.

## Results

### Survey of Conventional Stabilizing Agents

Water removal through lyophilization or freeze drying remains the standard strategy to stabilize RNA-LNP formulations by minimizing RNA and lipid hydrolysis^14,15,19,21,22,28,29^. As a cost-effective alternative, vacuum drying has been explored to preserve food^30^ or pharmaceutical materials^31^. However, its applicability to clinically used RNA-LNPs remains unexplored, with prior successes limited to pre-clinical LNP compositions (Fig. 1 a and Supplementary Table 1). In an attempt to stabilize clinically used LNPs, we formulated vacuum-dried Firefly Luciferase (FLuc) mRNA-LNPs and evaluated the *in vitro* transfection efficiency in A549 epithelial cells following reconstitution (Fig.1b). Freshly prepared LNPs in their respective buffer and excipient in solution served as positive controls. We synthesized six mRNA-LNP formulations with distinct combinations of ionizable and helper lipids (Fig.1b and d). Among these formulations, we included clinically used lipid compositions from Moderna’s mRNA-1273 (LNP A) and Pfizer/BioNTech’s BNT162b2 (LNP B) SARS-CoV-2 mRNA vaccines^1,2^, as well as the well-researched SORT (Selective ORgan Targeting) LNP (LNP F).^32,33^ To evaluate the impact of buffer composition on vacuum-dried LNP stability, we screened Tris and sodium acetate (NaAc) buffers, at pH 7.5 and 5.5, respective, as both buffers have demonstrated stabilizing effects on mRNA-LNPs in previous studies^5,20,34,35^.

**Fig. 1.**
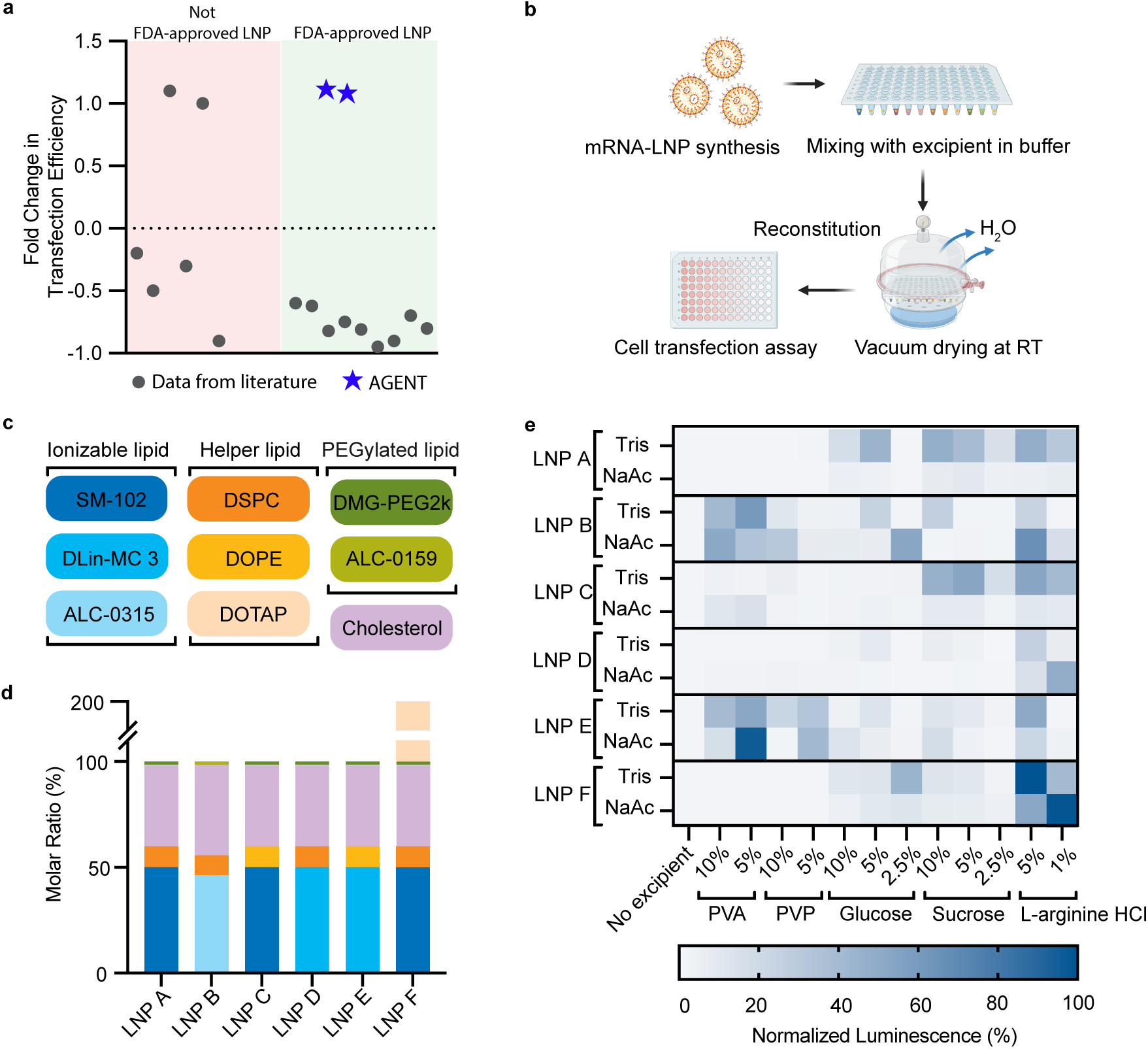
Exploration of the design space for solid-state mRNA-LNPs formulated via vacuum drying. **a** *In vitro* transfection efficiency of solid-state mRNA-LNPs normalized to initial soluble formulations from published studies employing vacuum-drying (Supplementary Table 1). **b** Schematic representation of the vacuum drying process for formulating and screening solid-state mRNA-LNPs. **c** Identity and **d** molar ratios of ionizable lipid, helper lipid, PEGylated lipid, and cholesterol used in the 6 LNP formulations for buffer and excipient screening. **e** Heatmap showing the *in vitro* transfection efficiency of reconstituted solid-state mRNA-LNPs in A549 cells for six different LNP compositions. Each composition was formulated sans excipient or using a single excipient at varying concentrations (w/v %) in either Tris or NaAc buffer. Luminescence values are normalized to the corresponding excipient with mRNA-LNPs maintained in solution.

As expected, we observed that vacuum-dried mRNA-LNPs in Tris or NaAc buffer alone showed almost 100% loss in transfection efficiency after reconstitution (Fig.1e), suggesting the need for a stabilizing agent. We therefore screened 40 excipients (Supplementary Fig. 1 and Fig. 1e), including sugars, amino acids, and water-soluble polymers, commonly employed in the pharmaceutical industry to stabilize biological materials ^36,37^. No single excipient was sufficient to stabilize mRNA-LNPs in the vacuum-drying process (Fig.1 e). Notably, the transfection efficiency of solid-state mRNA LNPs was strongly influenced by both buffer and excipient type in a composition-dependent manner, indicating that excipient compatibility varies across different LNP formulations. These results highlighted the necessity for tailored, LNP-specific excipient optimization rather than a one-size-fits-all approach. However, current studies on optimizing excipient formulations rely on one-variable-at-a-time experimentation,^14,15,19–22^ which is time-consuming and limits the ability to explore the entire design space. For example, even a modest 3-component screen of 40 excipients across 6 LNPs, with several concentration levels per component, would involve nearly 350,000 unique combinations, making exhaustive exploration impractical.

### Algorithm-guided Data-efficient Optimization Pipeline for Solid-state mRNA-LNPs

To accelerate the identification of optimal excipient formulations for solid-state mRNA-LNPs, we developed AGENT, a data-efficient experiment design pipeline based on Bayesian optimization algorithms. AGENT leverages prior experimental outcomes to iteratively refine its predictions, effectively balancing exploration of the design space with exploitation of high-performing formulations. This approach is particularly well-suited for problems where each experimental evaluation is time-and resource-intensive, enabling rapid optimization while minimizing the number of experiments ^38,39^. The implementation of AGENT unfolds in three stages: (1) constructing a Bayesian optimization algorithm with excipient types and concentration ranges as input variables^39^; (2) iteratively conducting *in vitro* screening based on excipient formulations suggested by the algorithm; and (3) validating the *in vivo* efficacy of solid-state mRNA-LNPs formulated with optimized excipient combinations. This framework is designed to efficiently converge on high-performing formulations within days, rather than the months of manual screening typically required.

In stage 1, we defined a five-dimensional design space to explore excipient combinations for solid-state mRNA-LNPs (Fig. 2a). The excipients were selected based on their established roles in stabilizing biologics, as well as their complementary physicochemical properties. Specifically, sucrose and glucose are widely used lyoprotectants and cryoprotectants that stabilize mRNA-LNPs ^40^. Polyvinyl alcohol (PVA) and polyvinylpyrrolidone (PVP) are water-soluble polymers that enhance matrix stability and provide the mechanical properties necessary for dissolvable MN fabrication ^18,23,24^.

**Fig. 2.**
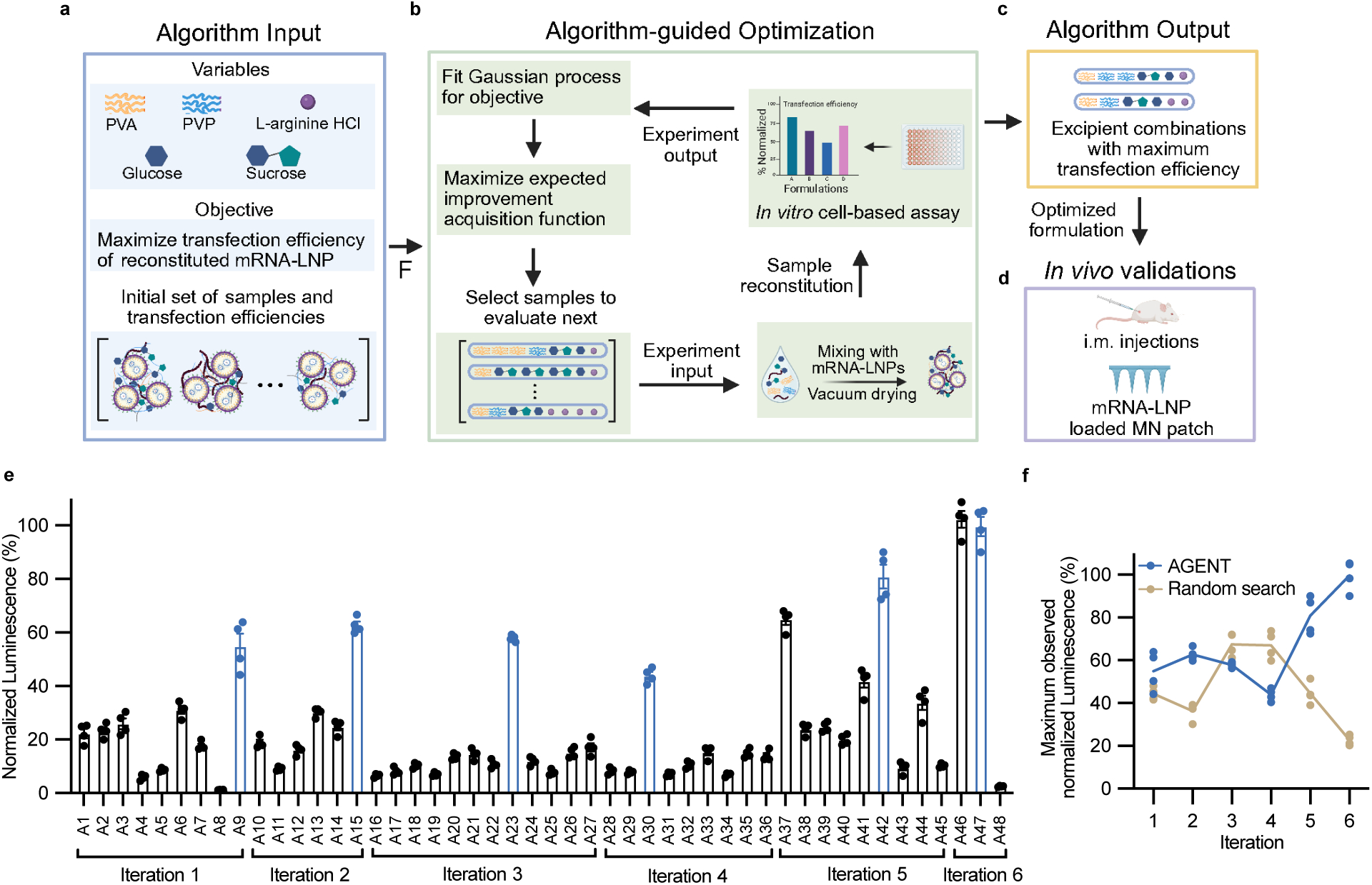
Overview of the algorithm-guided experiment pipeline for excipient optimization for solid-state mRNA-LNPs. **a** Required input information includes the variables and their ranges within the design space, the performance objective the algorithm aims to achieve, and an initial set of sample points to build an initial model. **b** Iterative process guided by the acquisition function to suggest new excipient combinations that are predicted to have better performance and add valuable information. Solid-state FLuc mRNA-LNPs are formulated with varying excipient combinations. In vitro transfection efficiency of reconstituted LNPs in A549 cells serves as input for the Bayesian optimization algorithm, which then proposes new excipient combinations for iterative testing. **c** Excipient combinations with the highest performance identified by the algorithm upon completing the optimization process. **d** Evaluation of *in vivo* efficacy for optimized solid-state mRNA-LNPs delivered via MN patches and i.m. injections. **e** Optimization progress of formulating solid-state Fluc mRNA-LNPs for LNP A. Best performing formulations from each iteration are highlighted in blue. **f** Optimization curves for 6-round search of the maximum luminescence using AGENT and random search. The same number of samples were tested per run for both methods.

Four excipients (PVP (MW 10,000), sucrose, glucose, and L-arginine) were treated as continuous variables ranging from 0 to 10% (w/v). For the fifth excipient (PVA, MW 31,000), single-excipient screening revealed that 5% PVA consistently outperformed 10% PVA across all LNP formulations (Fig. 1d). Therefore, we imposed a maximum concentration of 5% (w/v) on PVA. In the single-excipient screening experiments, Tris outperformed NaAc across most conditions (Fig. 1e), suggesting it provided a more favorable environment for preserving mRNA-LNP during vacuum drying. Therefore, to reduce the dimensionality of the Bayesian optimization search space, we fixed the buffer condition to Tris during AGENT optimization. Solid-state FLuc mRNA-LNPs were prepared by mixing the LNPs with selection excipient combinations, followed by vacuum drying. Transfection efficiency of reconstituted FLuc mRNA-LNPs was then measured *in vitro*. The transfection efficiencies from the initial set of randomly generated excipient combinations were used to train a Gaussian process surrogate model, which estimates both the predicted performance and associated uncertainty of new formulations. To choose the next formulation to test in each iteration, the algorithm utilized the Expected Improvement acquisition function, which quantifies the potential value of sampling a new point by balancing exploitation of discovered high-performing regions and exploration of unknown regions with high uncertainty^41^. The algorithm was implemented within AutODEx framework^39^.

In stage 2, the algorithm recommended new excipient combinations to test, and we iteratively repeated the cycle of formulation, drying, reconstitution, and in vitro assay. This process continued until a solid-state mRNA-LNP formulation achieved transfection efficiency comparable to its solution-state counterpart (Fig. 2b, c). In stage 3, performance of the optimized solid-state formulation obtained through *in vitro* experiments were then validated *in vivo* through i.m. injections after reconstituion or MN patch applications (Fig. 2 d).

### Performance of AGENT for Optimizing Solid-state FLuc mRNA-LNPs

We tested the proposed algorithm-guided experiment pipeline with FLuc mRNA encapsulated in LNP A and LNP B, Moderna and Pfizer formulations respectively, given their clinical relevance. We conducted 6 algorithm iterations for LNP A and 3 iterations for LNP B. To expedite the process, multiple formulations were tested in parallel during each iteration. In contrast to random search, which failed to identify solid-state formulations exceeding 80% normalized luminescence, Bayesian optimization learned from prior experiments and achieved an optimized formulation within 6 iterations (Fig. 2f). Optimized excipient formulations (sample ID A47 and B12) were achieved after testing a total of 48 and 12 samples for LNP A and B, respectively (Fig. 2e, Supplementary Fig. 2a). Solid-state mRNA-LNPs formulated with optimized excipient formulations achieved transfection efficiencies approximately 1.4-fold higher for LNP A and 1.3-fold higher for LNP B compared to the best-performing formulation from the initial randomly generated sample set in round 1. Additionally, there was a nearly 90% increase compared to manually optimized formulations (i.e., 5% sucrose, 5% PVA, and 5% sucrose with 5% PVA) (Fig. 3a, g).

**Fig. 3.**
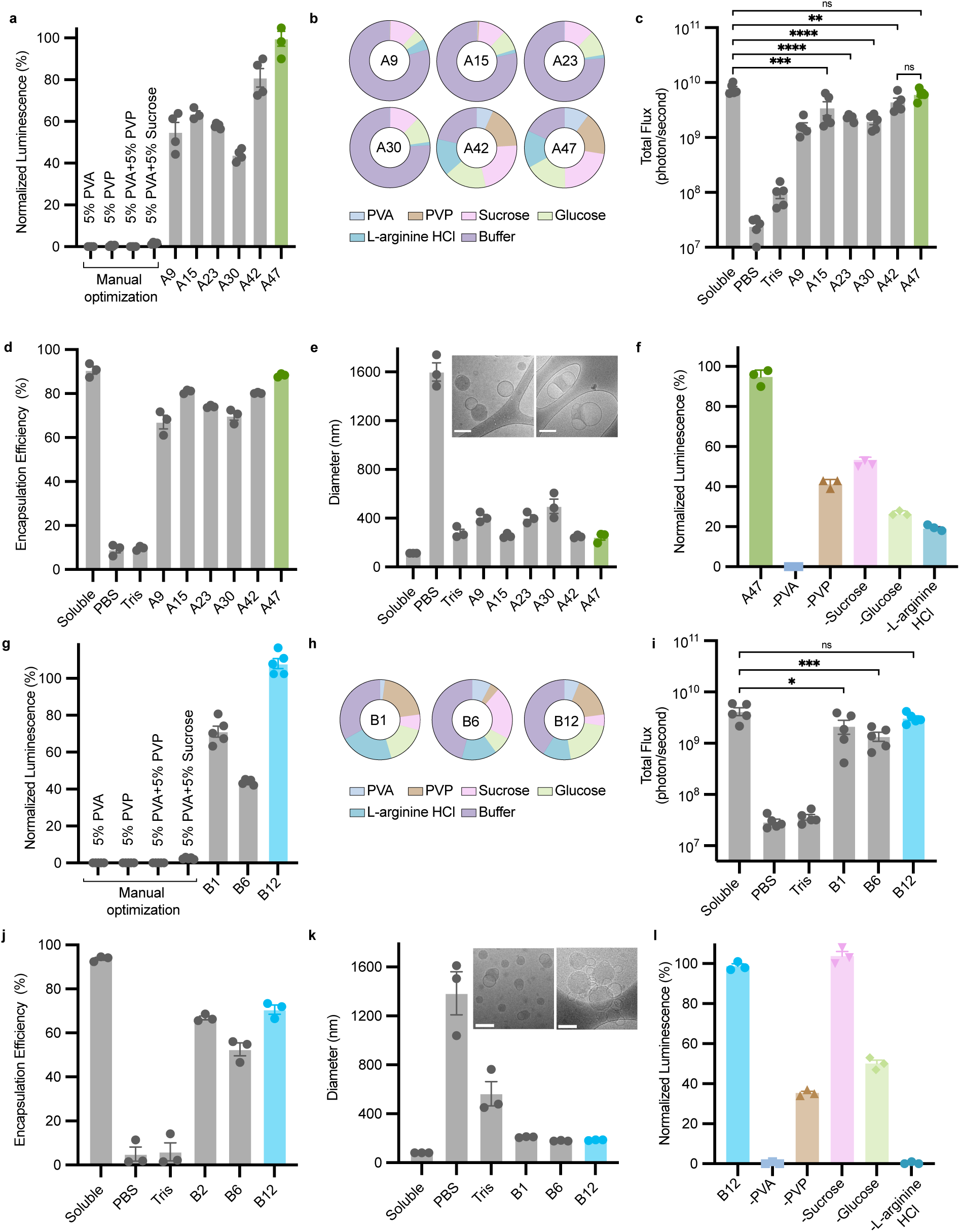
Optimizations, characterizations and *in vivo* validations of solid-state FLuc mRNA-LNPs. Highest *in vitro* transfection efficiency of solid-state FLuc mRNA-LNPs formulated with **(a)** LNP A and **(g)** LNP B lipid compositions from each optimization iterations. Solid-state mRNA-LNPs were formulated using excipients suggested by the algorithm, reconstituted in PBS, and treated with A549 cells. Luciferase expression was normalized against mRNA-LNPs with the same excipients maintained in solution. Optimized excipient formulations are highlighted in green for LNP 1 and in blue for LNP B. Composition % (w/v) of best performing excipient from each round of optimization for **(b)** LNP A and **(h)** LNP B. Quantification of luminescence intensity at the injection site in mice at 6 h following intramuscular administration of soluble and reconstituted solid-state FLuc mRNA-LNPs (0.3 μg mRNA per mouse) for **(c)** LNP A and **(i)** LNP B. n = 5 biologically independent mice per group, one-way ANOVA with Tukey’s test. mRNA Encapsulation efficiency of soluble and reconstituted solid-state FLuc mRNA-LNPs for **(d)** LNP A and **(j)** LNP B. Hydrodynamic diameter of soluble and reconstituted formulations measured by DLS for **(e)** LNP A and **(k)** LNP B. **Inset:** cryogenic transmission electron microscopy (cryo-TEM) image of soluble (left) and reconstituted solid-state FLuc mRNA-LNPs (right). Scale bars=100 nm. *In vitro* transfection efficiency of optimized solid-state FLuc mRNA-LNPs compared with formulations in which individual excipient components were removed for (f) LNP A and (l) LNP B.

To further validate the optimization process, we selected the top-performing formulation from each iteration, as well as solid-state mRNA-LNPs formulated with either PBS or Tris buffer to test *in vivo*. These formulations were reconstituted, administered intramuscularly into mice, and bioluminescence signals at the injection site were evaluated by IVIS. The IVIS data recapitulated the trends observed *in vitro*: the reconstituted optimized formulation showed protein expression levels comparable to soluble LNPs, while earlier iterations exhibited lower transfection efficiency (Fig. 3c, i). Furthermore, we formulated solid-state FLuc mRNA-LNPs with the optimized excipient combinations in both Tris and NaAc buffers (Supplementary Fig. 3). Consistent with the initial screening results, Tris-buffered formulations showed higher transfection efficiency than NaAc-buffered formulations. Solid-state mRNA-LNPs remained stable at 0°C to 45°C for 2 weeks (Supplementary Fig. 4a, b). Notably, they retained bioactivity after 1 year of storage at room temperature, whereas soluble LNPs stored at 4 °C showed an approximately 20-fold reduction in transfection efficiency (Supplementary Fig. 4c). The encapsulation efficiency of reconstituted optimized solid-state LNP A remained comparable to the freshly synthesized LNPs, while optimized solid-state LNP B showed a slight decrease in encapsulation efficiency (Fig. 3d, j). Additionally, the size of the reconstituted optimized mRNA-LNPs exhibited only minor changes. Notably, Tris-formulated solid-state mRNA-LNPs showed significantly less particle size increase compared to those formulated with PBS (Fig. 3e, f). Cryo-TEM images revealed that reconstituted solid-state mRNA-LNPs exhibited a core with multiple bleb-like structures rather than a single typical LNP morphology observed in soluble formulations (Fig. 3e, f). These structures might arise from lipid phase transitions and osmotic changes during drying-rehydrating process^42^. Gel electrophoresis analysis (Supplementary Fig. 5) showed that mRNA integrity was well preserved after vacuum drying, indicating that the excipients effectively protected the mRNA from degradation during solid-state formation. In parallel, quantification of oxidized ionizable lipid revealed oxidation levels below 0.0615% of total ionizable lipids after vacuum drying, supporting the chemical stability of the formulation (Supplementary Fig. 6). Furthermore, removal of individual excipient components from the optimized formulation resulted in a substantial decrease in transfection efficiency (Fig. 3f, l), except for sucrose in LNP B, likely due to its low proportion in the optimized formulation. These results highlight that the relative concentrations of individual excipients and their combinations are critical determinants of stability in solid-state mRNA-LNPs.

### Validation of Optimized Solid-state LNP Formulations Using RBD mRNA Vaccine Against SARS-CoV-2

Having established the performance of algorithm-optimized solid-state LNPs using FLuc mRNA as a model reporter, we next assessed their functional efficacy using mRNA encoding for the SARS-CoV-2 spike receptor-binding domain (RBD) protein. Various excipient conditions were tested for LNP A: no excipients, 5% PVA (w/v), 5% PVP (w/v), 5% PVA/5% sucrose (w/v), and the algorithm-optimized excipient formulation. Freshly prepared soluble mRNA-LNP and reconstituted solid-state samples were intramuscularly administered to mice, followed by a booster dose two weeks later. Mouse sera were collected weekly to quantify the antigen-specific antibody titer (Fig. 4a). The results show a strong correlation between vaccine induced antibody titer and excipients used to formulated solid-state mRNA-LNPs (Fig. 4b). Solid-state mRNA-LNP formulated with no excipients failed to generate a measurable IgG response in 4 out of 5 animals (Fig. 4b). LNPs formulated with 5% PVA elicited higher titers than those formulated with 5% PVP (Fig. 4b; P_PVA_=0.0046; P_PVP_=0.0002). The addition of 5% sucrose in 5% PVA increased the antibody response (Fig. 4b; P_PVA+sucrose_=0.0036). Algorithm optimized excipient formulations generated antibody titers that were not statistically distinguishable from the fresh soluble samples (Fig. 4b; p=0.9839). We observed the same outcome for solid-state LNP B-based RBD mRNA-LNPs using algorithm-generated excipient formulations specifically optimized for LNP B (Fig. 4c; p>0.9999). Furthermore, algorithm-optimized solid-state RBD mRNA-LNPs retained their efficacy after storage at room temperature and 37 °C for two months, eliciting antibody responses comparable to freshly reconstituted samples, thereby demonstrating excellent thermostability (Fig. 4d, e, Supplementary Fig. 6). To demonstrate the broader applicability of the AGENT-designed solid-state mRNA-LNP platform beyond SARS-CoV-2 vaccines, we expanded the study to include mRNA encoding for ovalbumin (OVA). Following reconstitution, solid-state formulations elicited strong immune responses comparable to freshly prepared soluble formulations (Supplementary Fig. 8), supporting the generalizability of the platform across different mRNA cargos and clinically relevant LNP systems.

**Fig. 4.**
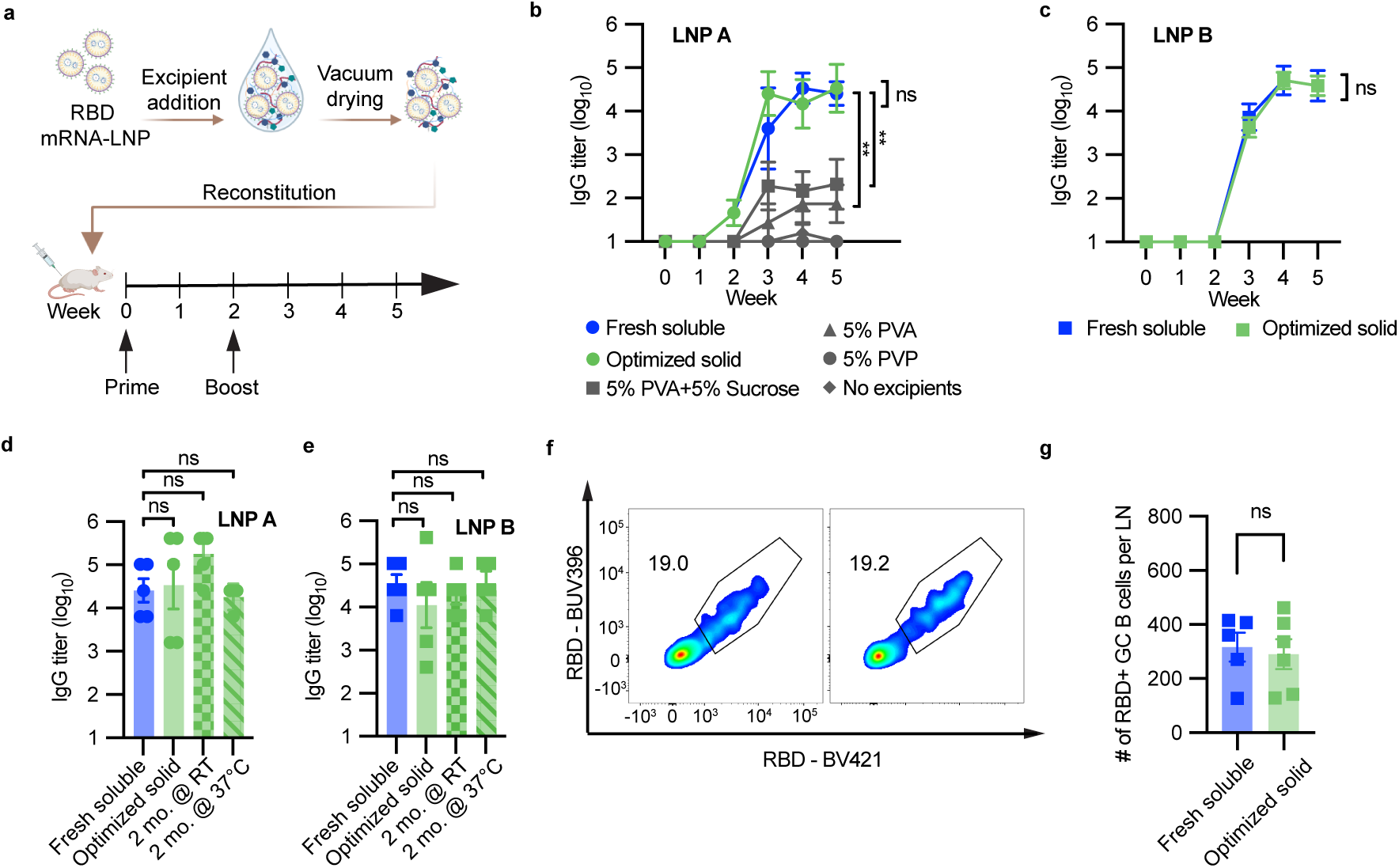
Optimized solid-state mRNA-LNP formulations for delivery of SARS-CoV-2 vaccine via injections. **a** Balb/c mice were immunized with soluble or reconstituted solid-state SARS-CoV-2 RBD mRNA-LNPs via i.m. injections followed by a booster dose 2 weeks later. Anti-RBD titers in serum were measured weekly. **b** Solid-state RBD mRNA-LNPs (LNP A lipid composition) were formulated with 5% PVA (w/v), 5% PVP (w/v), 5% PVA/5% sucrose (w/v), or the algorithm-optimized excipients. Plot shows serum anti-RBD protein IgG titers through week 5 (n=3 mice for 5% PVA and 5% PVP; n = 5 mice for other groups; 0.3 μg mRNA per mouse). **c** Mice were immunized with soluble or reconstituted solid-state SARS-CoV-2 RBD mRNA-LNPs via i.m. injections formulated with LNP B lipid composition (n = 5 mice; 0.3 μg mRNA per mouse). Plot shows serum anti-RBD protein IgG titers through week 5. Serum anti-RBD IgG titers at week 5 in mice administered fresh soluble or reconstituted solid-state RBD mRNA-LNP formulations with **d** LNP A and **e** LNP B lipid composition. Solid-state formulations were either fresh or stored at room temperature or 37°C for two months. **f** Mice were immunized i.m. with fresh soluble or reconstituted solid-state RBD mRNA-LNPs (LNP B lipid composition; n = 5 mice; 10 μg mRNA per mouse). Draining LNs were analyzed 10 days later. Representative flow cytometry plots of RBD-binding GC B cells. Fresh soluble (left) and reconstituted solid-state RBD mRNA-LNPs (right). **g** Cell count of RBD tetramer++GC B cells per LN. Statistical significance was determined by one-way ANOVA followed by Tukey’s multiple comparisons test [(b), (d), and (e)] or two-tailed unpaired *t* test [(c) and (g)].

In addition to assessing humoral immune responses, we evaluated germinal center (GC) responses induced by solid-state RBD mRNA-LNPs. BALB/c mice were immunized with fresh soluble or reconstituted optimized solid-state RBD mRNA-LNPs (10 μg/mouse) and draining lymph nodes were analyzed for GC B cells 10 days post immunization. The number of RBD-binding GC B cells, defined as B220+GL7+ RBD tetramer++cells (Supplementary Fig. 9), were comparable between two groups (Fig. 4f, g; p=0.7457), indicating that fresh soluble and solid-state LNPs recruited GC B cells to a similar extent *in vivo*. As GC activity mediates B-cell affinity maturation and the establishment of durable, high-quality antibody responses, these results demonstrate that solid-state LNPs can sustain functional vaccine efficacy comparable to that of liquid formulations.

### Delivery of Optimized Solid-state RBD mRNA Vaccine via MN patches in Rodents and Non-human Primates

MN patches represent a promising strategy for enhancing the accessibility of vaccines in low-resource settings by enabling self-application, eliminating sharps waste, reducing pain, and providing long-term shelf stability. Using the algorithm optimized excipient formulations, we fabricated RBD mRNA loaded MN patches for both LNP A and LNP B. Following a prime-boost dosing regimen, we vaccinated mice with RBD mRNA-LNP loaded MN patches at weeks 0 and 2 (Fig. 5a). ELISA results showed RBD mRNA-LNP loaded MN patches generated high antigen-specific IgG titers, comparable to those received intramuscular administration of fresh soluble mRNA-LNPs or reconstituted solid-state mRNA-LNPs with optimized excipients (Fig. 5b). To assess neutralizing activity, we performed an RBD-angiotensin converting enzyme 2 (ACE2) binding assay on serum collected in week 4. Neutralizing antibody levels were comparable across groups, indicating that optimized solid-state RBD mRNA-LNPs delivered through injections or MN patches provided protection equivalent to intramuscular injection of fresh soluble LNPs (Fig. 5c).

**Fig. 5.**
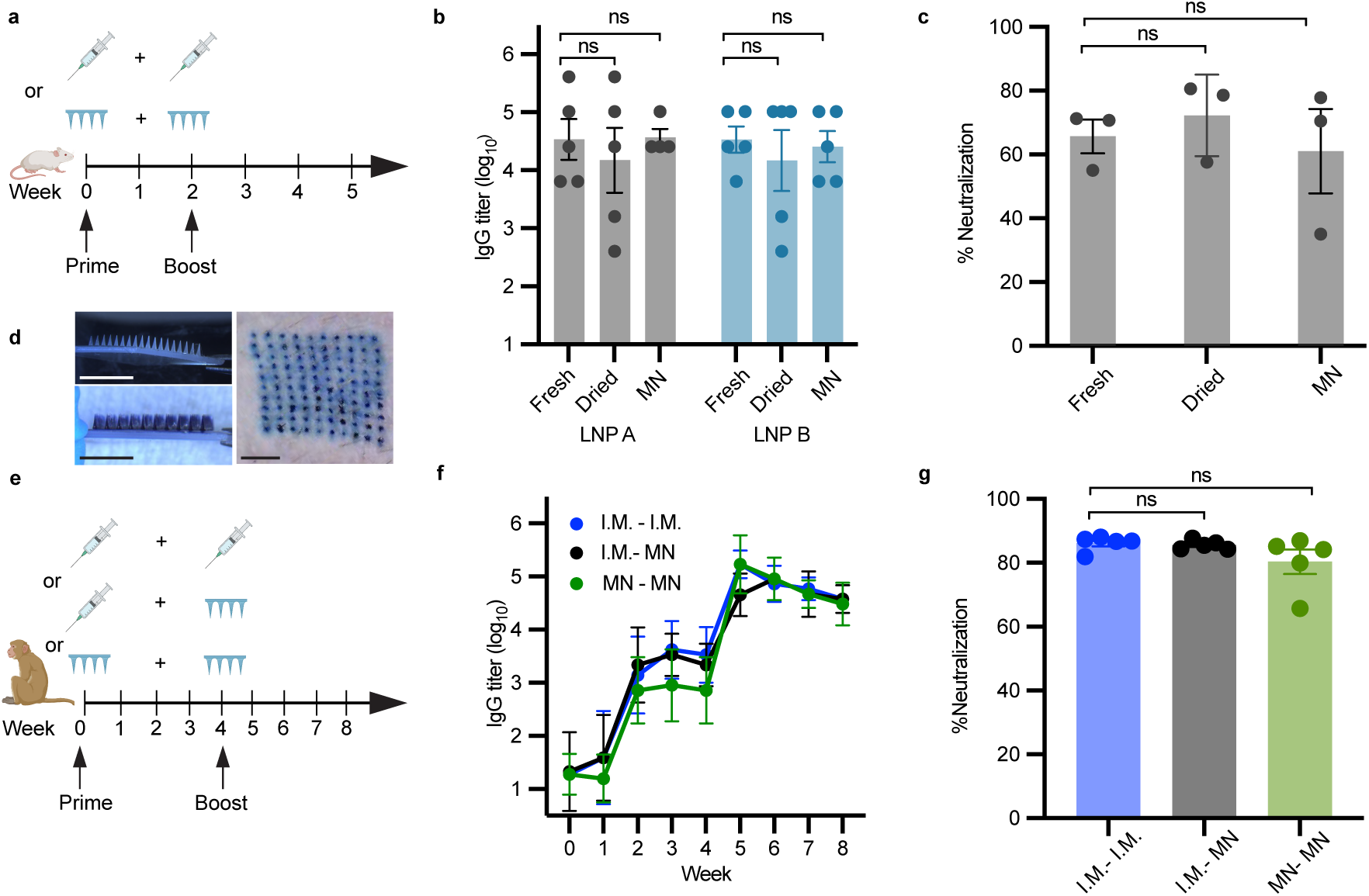
Optimized solid-state mRNA-LNP formulations for delivery of SARS-CoV-2 vaccine via MN patches in mice and NHPs. **a** Balb/c mice were immunized with soluble or reconstituted solid-state SARS-CoV-2 RBD mRNA-LNPs via i.m. injections or SARS-CoV-2 RBD mRNA-LNP loaded MN patches followed by a booster dose 2 weeks later (n = 5 mice; 0.3 μg mRNA per mouse per dose). **b** Quantification of serum anti-RBD protein IgG titers at week 4. **c** Percent neutralization of SARS-CoV-2 antibodies determined by RBD-ACE2 binding assay, using serum collected in week 4 at 1:50 dilution for LNP A. **d** Representative images of RBD mRNA-LNP loaded (left top) and trypan blue loaded (left bottom) MN patch fabricated with optimized excipient formulation. Optical image of trypan blue staining, showing puncture marks caused by the penetration of MNs on NHP skin. Scale bar = 7.5 mm. **e** Schematic diagram of vaccination in cynomolgus macaques. Cynomolgus macaques were immunized i.m. or with MN patches with 30 μg of RBD mRNA-LNPs (LNP B lipid composition) and boosted with the same dose on day 28 after first vaccination. **f** SARS-CoV-2 RBD-specific IgG antibody titers of the immunized animals were determined by ELISA. **g** RBD-ACE2 binding assay was performed to determine % neutralization of SARS-CoV-2 antibodies in cynomolgus macaques, using serum collected in week 8 at 1:50 dilution. Statistical significance was determined by one-way ANOVA followed by Tukey’s multiple comparisons test.

As a step towards the clinical translation of MN patches fabricated with algorithm-optimized solid-state RBD mRNA-LNPs, we vaccinated cynomolgus macaques with MN patches and compared the immune responses to intramuscular injections. As a pilot study, we first tested skin penetration using a trypan blue-loaded MN patch, which confirmed that MNs incorporating our optimized excipients provided sufficient mechanical strength for successful skin penetration in NHPs (Fig. 5d). Using LNP B-based RBD-mRNA LNPs, we vaccinated groups of cynomolgus macaques at weeks 0 and 4 with three dosing strategies: i.m. prime and boost (i.m. - i.m.), i.m. prime with a MN boost (i.m. - MN), or MN prime and boost (MN - MN) (Fig. 5e). All three groups elicited strong humoral responses across 8 weeks of the study (Fig. 5f and g), demonstrating that AGENT enables MN patches to serve as an effective and practical alternative mRNA vaccination strategy to intramuscular injection.

## Discussion

Cold-chain requirements add substantial infrastructure costs throughout the product life cycle, including specialized packaging, ultra-cold storage, temperature-controlled transport, and continuous monitoring. Compared with cold-chain vaccines, thermostable vaccine formulations could reduce storage costs by an estimated 71-86% and decrease wastage costs by 50% or more ^43,44^. Developing solid-state mRNA-LNP formulations is a key step toward improving the stability and accessibility of mRNA vaccines. While lyophilization or freeze-drying can extend mRNA-LNP shelf life, it is resource-intensive^15,19^ and unsuitable for advanced delivery systems such as MN patches. Recent studies have explored vacuum drying, but progress has been constrained by narrow excipient screening and inefficient single-variable testing^18,23,24,35^. Importantly, results from previous studies and this study, show that LNP composition critically influences mRNA-LNP recovery after drying ^23,24^, highlighting the need for strategies that account for LNP formulation diversity.

While AI-driven and deep learning approaches are increasingly being applied to accelerate formulation and drug-delivery development, many methods remain constrained by the need for large, high-quality training datasets. In our case, the optimization of vacuum-dried mRNA-LNP formulations represents a highly specialized and underexplored experimental space, for which no public datasets exist, and data generation is inherently costly. Under these constraints, Bayesian optimization is more appropriate, as it is specifically designed for optimization in small-sample regimes. Bayesian optimization has been widely adopted in materials discovery, such as nanoparticle synthesis^45^ and 3D printing material development^46^, to our knowledge, this work represents its first application to stabilizing mRNA-LNP vaccines. This versatile approach can be readily applied to stabilize different biologics, including proteins, antibodies, nanocarriers. Our algorithm-guided screening pipeline enables high-throughput, data-efficient identification of stabilizing excipient combinations. This approach is not only more efficient than traditional one-variable-at-a-time screening but is also broadly applicable across different LNP compositions.

Here, we demonstrate the successful optimization of excipients for two clinically relevant LNP formulations, highlighting the versatility and robustness of our strategy. Stabilization of certain LNP formulations, such as the Moderna-like LNP (LNP A), has proven particularly challenging in prior work, with improvements typically achieved only through modification of the LNP composition itself ^21,23,24^. However, altering the lipid composition introduces uncertainty regarding immune responses and safety ^47–50^, and preserving clinically approved LNP formulations is advantageous for translational purposes. Our findings show that incorporating suitable combinations of widely used excipients can effectively stabilize diverse LNPs in a solid state without requiring changes to the lipid composition. This provides a flexible library of excipient formulations ideally suited for downstream applications, including the fabrication of MN patches.

In our study, bleb-like structures were observed in reconstituted solid-state mRNA-LNP formulations, suggesting that the drying and reconstitution process potentially promoted this morphology. Several reports suggest that bleb-like morphologies may enhance LNP stability and efficacy^42,51^. Nevertheless, the presence of bleb structures warrants consideration from both regulatory and development perspectives. The bleb structures are not currently specified among the conventional critical quality attributes (CQAs) for LNP products which include particle size, size distribution, polydispersity index, encapsulation efficiency, and mRNA integrity ^52^. While morphology and lipid organization offer valuable insights into LNP function, such structural features are not yet classified as CQAs due to measurement challenges, their heterogeneous nature, and the lack of generalizable rules to derive structure-efficacy relations ^52^. Addressing this gap will require well-controlled manufacturing parameters and development of standardized analytical methods beyond cryo-EM, including advanced imaging, spectroscopic, and quantification techniques ^53^. Systematic optimization of formulation and processing conditions is recommended to achieve reproducible bleb formations and ensure consistent *in vivo* performance, product uniformity, and batch-to-batch reproducibility.

Finally, this study provides, to our knowledge, the first demonstration of successful delivery of mRNA vaccines in non-human primates using MN patches. These results establish MN patches as a promising platform for mRNA vaccine administration, supporting their feasibility for human use. More broadly, by enabling solid-state stabilization, this approach could reduce reliance on cold-chain infrastructure, which is a major contributor to vaccine cost and wastage. Because our vacuum-drying process is a one-time manufacturing step and uses established, FDA-approved materials already common in pharmaceutical products, we do not anticipate major added material costs. Taken together, our work introduces a broadly applicable stabilization strategy and highlights the potential of combining solid-state mRNA-LNPs with MN technology to enable accessible, thermostable, and user-friendly vaccine delivery.

## Acknowledgments

We thank the Koch Institute’s Robert A. Swanson (1969) Biotechnology Center (RRID:SCR_018674) for technical support, specifically the Nanotechnology Materials core, the BioMicroCenter, and the Animal Imaging and Preclinical Testing facilities. We thank Dr. Namit Chaudhary for his guidance on tissue harvesting and flow cytometry. We thank Pavle Konaković and Yunsheng Tian for their help with algorithm implementation and use of AutODEx.

