## Supplemental Files for "Accelerated discovery of thermostable vaccines using data-efficient AI"

#### Methods

##### mRNA-LNP synthesis

Cholesterol, 1,2-dimyristoyl-rac-glycero-3-methoxypolyethylene glycol-2000 (DMG-PEG 2000), 1,2-dioleoyl-sn-glycero-3-phosphoethanolamine (DOPE), and 1,2-distearoyl-sn-glycero-3-phosphocholine (DSPC) were purchased from Avanti Polar Lipids. SM-102, D-Lin-MC3-DMA (MC3), ALC-0315, and ALC-0159 were obtained from MedChemExpress. Purified CleanCap mRNA constructs encoding for Firefly Luciferase (FLuc), SARS-CoV-2 Spike Protein (RBD) or Ovalbumin (OVA) were purchased from TriLink Biotechnologies, stored at -80°C before use. Lipid nanoparticles (LNPs) were synthesized using an organic-aqueous precipitation method in an automated microfluidic mixer. The organic phase was prepared by dissolving lipids in ethanol at molar ratios of 50:10:38.5:1.5 (SM-102 or MC3: DOPE or DSPC: cholesterol: DMG-PEG 2000) or 46.3:9.4:42.7:1.6 (ALC-0315: DSPC: cholesterol: ALC-0159). The aqueous phase contains mRNA in 10 mM citrate buffer at pH 3.0. The two phases were prepared at a volume ratio of 3:1 and mixed with a NanoAssemblr Ignite instrument (Precision Nanosystems, Vancouver, Canada) at a 12 mL/min flow rate. The collected LNPs were dialyzed against pH 7.4 PBS at 4 °C for 1 hour using a 10,000 molecular weight cutoff (MWCO) cassette followed by another 2-hour dialysis in DI water at 4 °C. mRNA-LNPs were then concentrated fourfold by centrifuging at 3,000g in an Amicon filter. LNPs were stored at 4 °C before use.

##### Solid-state mRNA-LNP formulation with excipients

Stock excipient solution of polyvinyl alcohol (PVA, Mw 31,000), polyvinylpyrrolidone (PVP, Mw 10,000), sucrose, glucose, L-arginine hydrochloride was prepared with 20 mM RNase-free Tris buffer (pH 7.5) or sodium acetate buffer (pH 5.5). Excipient solutions were then mixed at predetermined weight to volume ratios. Solid-state mRNA-LNPs were formulated by combining mRNA-LNPs with excipient solution at a 1:5 (mRNA:excipients, w/v) ratio. Aliquots of 15 µL was then dispensed into an Eppendorf tube and dried in a vacuum chamber containing silica gel desiccant, connected to a vacuum pump maintained at approximately -60kPa. Samples were dried under ambient temperature for 72 hours to ensure complete water removal. Solid-state mRNA-LNPs were reconstituted in 1x PBS buffer at 0.03 µg/µL mRNA concentration.

##### mRNA-LNP characterization

The size of LNPs was measured using a Malvern ZetaSizer Nano-NS (Malvern Instruments) following a 200-fold dilution in ultra-pure water. Quan-iT RiboGreen RNA Assay Kit (Thermo Fisher Scientific) was used to quantify the mRNA concentration and encapsulation efficiency of LNPs. Briefly, LNPs were diluted in equal volume of Tris-EDTA buffer (TE) or 2% Triton X-100 in Tris-EDTA buffer (TX). RiboGreen reagent was then added to each sample and incubated for 15 min at 37 °C. The fluorescence intensity (excitation/emission: 485/515 nm) was measured on a Tecan Spark Multimode Microplate reader. The encapsulation efficiency of LNPs was calculated by $\frac{TX-TE}{TX} * 100\%$.

**LNP Oxidation Assay**

The concentration of oxidized SM102 in dried samples was determined using the OxiSelect™ Total Antioxidant Capacity colorimetric assay (Cell Biolabs inc., San Diego, CA) by employing a protocol modified from the manufacturer. LNPs were prepared and vacuum dried as previously described. Samples were rehydrated in 1:1 $H_2O$:MeOH at a final SM-102 concentration of ~2.4 mM. Positive control LNPs were prepared by replacing the ionizable lipid SM-102 in the LNP formulation with SM-102 N-oxide (Cayman Chemical, Ann Arbor, MI) to mimic LNPs that had completely oxidized. All reagents in the OxiSelect™ kit were prepared in MeOH. To perform the assay, 60 µL of standards and samples were added to 90 µL of 2X reaction buffer and 50 µL of 1X Cu ion reagent in a clear 96-well plate. The reaction was allowed to proceed for 5 min at RT and MeOH-insoluble excipients that precipitated during the incubation period were removed by centrifugation. The final absorbance at 490 nm was measured using a microplate reader (Tecan Life Sciences, Männedorf, Switzerland). The assay was validated using an $H_2O_2$ standard curve. The concentration of oxidized SM-102 in the samples was determined by preparing a standard curve of SM-102 N-oxide with a dynamic range of 4 mM-0.0625 mM.

**Cryogenic-TEM**

RBD mRNA-LNPs were used for all samples. Soluble mRNA-LNPs were diluted in ultra-pure water and solid-state mRNA-LNPs were reconstituted in in ultra-pure water. Samples were plunge-frozen using a 930 Gatan Cryoplunge 3. All the samples were imaged using a JEOL 2100 FEG microscope at 200-kV acceleration voltage.

##### *In vitro* high-throughput screening

For algorithm guided *in vitro* optimization, 5,000 A549 cells per well were seeded in white 96-well plates. After overnight incubation, reconstituted solid-state FLuc mRNA-LNPs and soluble FLuc mRNA-LNPs with excipients were added at 50 ng per well. Luminescence was measured 24 h after transfection using Bright-Glo (Promega) according to the manufacturer's instructions. Luminescence was quantified using a Tecan Infinite M200 Pro plate reader (Tecan).

##### Animal studies

All animal experiments were authorized by the Division of Comparative Medicine at the Massachusetts Institute of Technology (#2208000412). Six-week-old female BALB/c mice were purchased from Charles River Laboratory and housed in an MIT animal facility. For nonhuman primate experiments, fifteen (five females, ten males) genetically unselected cynomolgus macaques ranging in age from 1 year 2 months to 6 years 4 months were purchased from and housed at Alpha Genesis, Inc (Yemassee, SC).

##### *In vivo* mouse imaging

To qualitatively assess the efficacy of solid-state mRNA-LNPs compared with soluble mRNA-LNPs, we intramuscularly (i.m.) injected 0.3 µg mRNA doses of the LNPs loaded with FLuc in the right quadricep muscle of BALB/c mice. At 6 h following injections, mice were injected intraperitoneally with 150 $\mu$L of d-luciferin substrate (30 mg/mL). Mice were anesthetized in a ventilated anesthesia chamber with 1.5% isofluorane in oxygen. 15 min after d-luciferin injection, luminescence was measured with an in vivo imaging system (IVIS, PerkinElmer) and quantified using LivingImage software (PerkinElmer).

##### Mouse Vaccination

To assess the efficacy of solid-state mRNA-LNPs and compared with soluble mRNA-LNPs using mRNA encoding for the SARS-Co-V-2 receptor binding domain, we vaccinated healthy Balb/C mice by injecting 0.3 µg mRNA doses of the LNP-RNA i.m. in the right quadricep muscle. For RBD mRNA-LNP loaded MN patch applications, two 4 x 4 MN patches were applied to the left and right footpad of mice with anesthesia, with a 10-min wear time for each, for a total dose of 0.3 µg mRNA per mouse. For all experiment groups, mice were boosted with 0.3 µg RBD mRNA via the same administration method (i.m. or MN patches) on day 14 after first vaccination. The mice underwent submandibular bleeding weekly. Serum was isolated by centrifuging blood at 10,000 ×*g* for 3 min and stored at −20 °C prior to use.

**Macaque Vaccination**

Healthy cynomolgus macaques were randomly assigned to three groups (n=5 per group) for immunization with RBD mRNA LNPs. The first group received a prime i.m. injection (30 µg mRNA) followed by a microneedle patch boost (30 µg mRNA) on day 28. The second group received both prime and boost via MN patches. A third group, serving as the positive control, was immunized via i.m. injection for both prime and boost. Blood samples were collected prior to the initial vaccination and weekly thereafter. Serum was analyzed for SARS-CoV-2 RBD-specific IgG titers and neutralizing antibody levels.

**Anti-OVA and SARS-CoV-2 anti-RBD titer quantification**

To quantify SARS-CoV-2 RBD-specific antibodies, enzyme-linked immunosorbent assays (ELISAs) were performed using clear high-binding 96-well plates (Corning). Plates were coated overnight at 4 °C with purified SARS-CoV-2 Spike RBD-His recombinant protein (SinoBiological): 1 µg/mL for mouse serum and 0.1 µg/mL for NHP serum. For anti-OVA IgG tier quantification, plates were coated with OVA protein (InvivoGen, vac-stova) at 20 µg/mL. Plates were then blocked for 2 hours at 37°C with BLOTTO blocking buffer (5% nonfat dry milk in 1× PBS with 0.05% Tween-20). Serum samples were incubated at 37°C for 2 hours, followed by detection using goat anti-mouse IgG-HRP conjugated antibody (SouthernBiotech, 1030-05), diluted 1:10,000 for mouse serum and goat anti-monkey IgG-HRP conjugated antibody (abcam, ab112767), diluted 1:15,000 for NHP serum. The reaction was developed using 3,3′,5,5′-tetramethylbenzidine (TMB) substrate, and the absorbance was measured at 450 nm with background subtraction at 630 nm using a microplate reader. Endpoint titers were defined as the highest dilution of serum that produced an optical density (OD) value at least three times greater than the background signal from pre-vaccination serum samples.

**Flow cytometry**

All staining steps were carried out at 4°C in FACS buffer (1x PBS with 2% heat inactivated FBS and 2 mM EDTA). Single cell suspensions were Fc blocked with anti-CD16/CD32 monoclonal antibody (mAb) and stained with Fixable Viability dye eFluor780. Cells were washed and incubated for 30 minutes with a cocktail of fluorescently labeled anti-mouse mAbs containing CD38-BB515, GL7-APC, PD1-BV605, CD4-BV711, CXCR5-PE, B220-PE-Cy7 and biotinylated BUV 396-conjugated and BV421-conjugated Spike RBD-His recombinant protein. Next, cells were washed and fixed with 0.5% paraformaldehyde (PFA) for 30 minutes. To obtain accurate cell counts, precision count beads (BioLegend) were added to each sample. All samples were acquired on a 5 laser FACSymphony A3 (Becton Dickinson) and data analyzed in FlowJo v10 (Treestar).

**mRNA-LNP loaded microneedle (MN) patch fabrication**

MN patches were fabricated in a two-step molding process on PDMS molds (Sylgard 184, Dow Corning) as previously reported [1]. Each MN patch comprised an array of 4 x 4 microneedles (1500 $\mu$m in length with a 400 µm x 400 µm base). Mixture of excipients and mRNA-LNPs was dispensed on the PDMS mold. For LNP A, the optimized excipient composition was 4.5326:7.8501:10:7.741:6.7613 (PVA : PVP : sucrose : glucose : L-arginine; w/v%). For LNP B, the optimized composition was 2.7659:7.5269:2.102:8.9558:5.1183 (PVA:PVP:sucrose:glucose:L-arginine; w/v%). Vacuum was applied at -90 kPa gauge pressure underneath the PDMS mold. MN patches were dried in a vacuum chamber (-60kPa gauge pressure) with desiccant for three days and demolded with double-sided tape.

**Bayesian optimization algorithm**

The optimization began with a real-valued objective function defined over $\mathcal{X}$, a domain of potential excipient combinations. Let $f: \mathcal{X} \rightarrow R$ denote the *in vitro* transfection efficiency, the objective function. The goal of optimization was to search this domain to identify a formulation $x^*$ that achieved the global maximum of the objective function $f^*$:

$$x^* \in \arg\max_{x \in \mathcal{X}} f(x); \quad f^* = \max_{x \in \mathcal{X}} f(x) = f(x^*)$$

$y = f(x) + \varepsilon$ described the measured transfection efficiency from formulation $x$, where $\epsilon$ represented measurement errors. The algorithm utilized Gaussian Process (GP) as the surrogate model of the formulation space:

$$p(f) = \mathcal{GP}(f; \mu, K)$$

Where the mean function $\mu(x) = E[\phi|x]$ determined the expected function value $\phi = f(x)$ at any location $x$, and the covariance function $K(x, x') = \text{cov}[\phi, \phi' \mid x, x']$ determined how deviations from the mean were structured. In this case, we use zero mean function and a Matérn

1/2 kernel: $K_{M1/2}(x, x') = \sigma_f^2 \exp\left(-\sum_{i=1}^{d} \frac{|x_i - x_i'|}{l_i}\right)$. The corresponding hyperparameters are specified in Supplementary Table 2.

To initialize the GP surrogate model, *in vitro* transfection efficiency from 13 formulations (9 randomly generated samples uniformly spanning the design space and 4 manually selected primary formulations) were used. Extending the GP on $f$ to include the entries of observed data yielded $p(f, y) = \mathcal{GP}\left(\begin{bmatrix} f \\ y \end{bmatrix}; \begin{bmatrix} \mu \\ m \end{bmatrix}, \begin{bmatrix} K & \kappa^T \\ \kappa & C \end{bmatrix}\right)$. [2] Writing $\mathcal{D} = y$ for the observed data, the GP posterior of $f$ could be derived as $p(f \mid \mathcal{D}) = \mathcal{GP}(f; \mu_{\mathcal{D}}, K_{\mathcal{D}})$, where

$$\mu_{\mathcal{D}}(x) = \mu(x) + \kappa(x)^T C^{-1}(y - m);$$

$$K_{\mathcal{D}}(x, x') = K(x, x') - \kappa(x)^T C^{-1} \kappa(x')$$ [2]

Given the probabilistic GP posterior, the expected improvement acquisition function (EI), assigns a score to each candidate formulation $x$ based on the expected benefit of sampling there:

$$\alpha_{\mathrm{EI}}(x; \mathcal{D}) = \int [max\ \mu_{\mathcal{D}'}(x')]\, p(y \mid x, \mathcal{D}) - max\ \mu_{\mathcal{D}}(x)$$ [3]

In each iteration, maximizing expected improvement will select points with predicted maximal value. The algorithm proposed a batch of formulations with batch size ranging from 3 to 9 samples for parallel evaluation to reduce experimental time. Newly acquired data were incorporated into the training set, the model was refitted on the new set, and the optimization cycle was repeated until the objective was achieved. The procedure described above was implemented within AutODEx framework[4]. Additional implementation details can be found at autodex.ai.

**Statistical methods**

Statistical analyses were conducted using GraphPad Prism software. Significance levels were represented by ns: no significance; *p ≤ 0.05; *p ≤ 0.01; ***p ≤ 0.001; ****p≤ 0.0001.

#### SUPPLEMENTARY Table

**Supplementary Table 1. Change in *in vitro* transfection efficiency of solid-state mRNA-LNPs after vacuum drying, normalized to corresponding soluble formulation.**

| LNP compositions | IL/HL/Chol/PEG | Clinically used | Fold change | Excipient(s) | Ref. |
|---|---|---|---|---|---|
| SM102/DSPC/Chol/DMG-PEG2000 | 50/10/38.5/1.5 | Yes | -0.6 | 10% PVA+10% sucrose | 24 |
| SM102/DSPC/Chol/DMG-PEG2000 | 60/10/38.5/1.5 | No | -0.2 | 10% PVA+10% sucrose | |
| SM102/DSPC/Chol/DMG-PEG2000 | 65/10/38.5/1.5 | No | -0.5 | 10% PVA+10% sucrose | |
| SM102/DSPC/Chol/DMG-PEG2000 | 50/10/38.5/1.5 | Yes | -0.62 | 5% PVA | 23 |
| SM102/DSPC/Chol/DMG-PEG2000 | 50/10/38.5/1.5 | Yes | -0.82 | 5% PVA+2.5% sucrose | |
| SM102/DSPC/Chol/DMG-PEG2000 | 50/10/38.5/1.5 | Yes | -0.75 | 5% PVA+10% sucrose | |
| SM102/DSPC/Chol/DMG-PEG2000 | 50/10/38.5/1.5 | Yes | -0.81 | 5% PVA+15% sucrose | |
| SM102/DSPC/Chol/DMG-PEG2000 | 50/10/38.5/1.5 | Yes | -0.95 | 5% PVA+10% trehalose | |
| SM102/DSPC/Chol/DMG-PEG2000 | 50/10/38.5/1.5 | Yes | -0.9 | 5%PVA+10% glucose | |
| SM102/DSPC/Chol/DMG-PEG2000 | 50/10/38.5/1.5 | Yes | -0.7 | 5% PVA +10% sucrose/ glucose | |
| SM102/DSPC/Chol/DMG-PEG2000 | 50/10/38.5/1.5 | Yes | -0.8 | 5% PVA+10% sucrose/trehalose/glucose | |
| Lipid5/DOPE/Chol/C14-PEG2000 | 50/10/38.4/1.5 | No | 1.1 | 20% PVA/PVP | 18 |
| Lipid5/DOPE/Chol/C14-PEG2000 | 50/10/38.4/1.5 | No | -0.3 | 20%PVP | |
| Lipid5/DOPE/Chol/C14-PEG2000 | 50/10/38.4/1.5 | No | 1.0 | 20% PVA/sucrose | |
| Lipid5/DOPE/Chol/C14-PEG2000 | 50/10/38.4/1.5 | No | -0.9 | 20% PVP/sucrose | |

| | | | | | |
|---|---|---|---|---|---|
| ALC0315/DSPC/Chol /ALC-0159 | 46.3/9.4/42.7/1.6 | Yes | 1.08 | 6.15%PVA+16.73%PVP+ 4.67%sucrose+19.9%glucose+ 11.37%L-arginine HCl | This work |
| SM102/DSPC/Chol/D MG-PEG2000 | 50/10/38.4/1.5 | Yes | 1.03 | 10.07%PVA+17.45%PVP+ 22.22%sucrose+17.2%glucose +15.03%L-arginine HCl | |

C14-PEG2000: 1-2-dimyristoyl-sn-glycero-3-phosphoethanolamine-N-[methoxy(polyethylene glycol)-2000] (ammonium salt)

Lipid 5: heptadecane-9-yl 8-((2-hydroxyethyl)(8-nonyloxy)-8-oxoctyl)amino)octanoate

**Supplementary Table 2. Gaussian Process hyperparameters used in optimization algorithm.**

| Hyperparameter | Initial Value | Optimization Range |
|---|---|---|
| $l$ | $(1, \ldots, 1) \in R^d$ | $\left(\sqrt{10^{-3}}, \sqrt{10^3}\right)$ |
| $\sigma_f$ | 1 | $\left(\sqrt{10^{-3}}, \sqrt{10^3}\right)$ |
| $\sigma_n$ | $10^{-2}$ | $(e^{-6}, 1)$ |

### SUPPLEMENTARY FIGURES

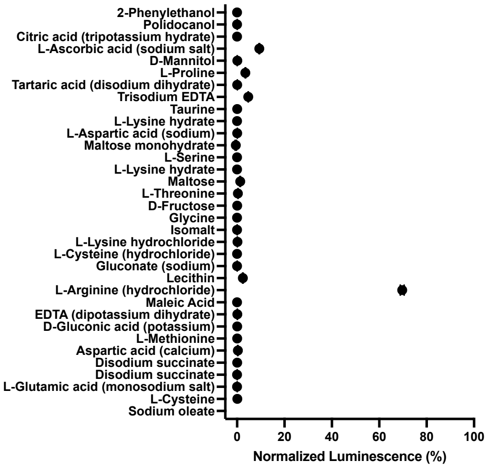


**Supplementary Fig. 1** ***In vitro*** **screening of amino acids for stabilizing mRNA-LNPs in solid states.** Solid-state mRNA-LNPs (LNP A) were formulated with 10% (w/v) amino acids dissolved in 20 mM Tris buffer via vacuum drying, reconstituted in PBS, and treated with A549 cells. Luciferase expression was normalized against mRNA-LNPs with the same excipients maintained in solution.

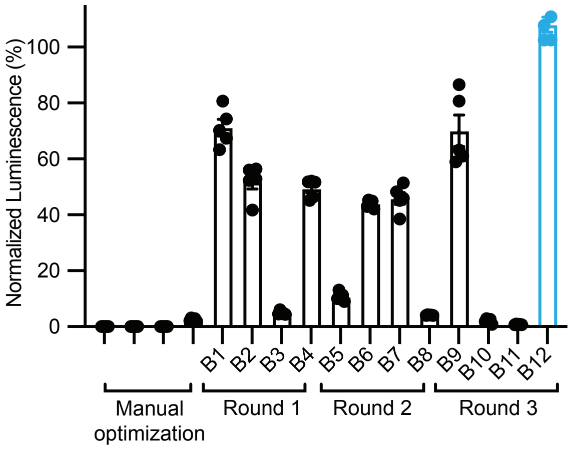


**Supplementary Fig. 2 Optimization progress of formulating solid-state Fluc mRNA-LNPs.** Plot showing improvement of *in vitro* transfection efficiency of solid-state Fluc mRNA-LNPs for LNP B. Optimized excipient formulation is highlighted in blue.

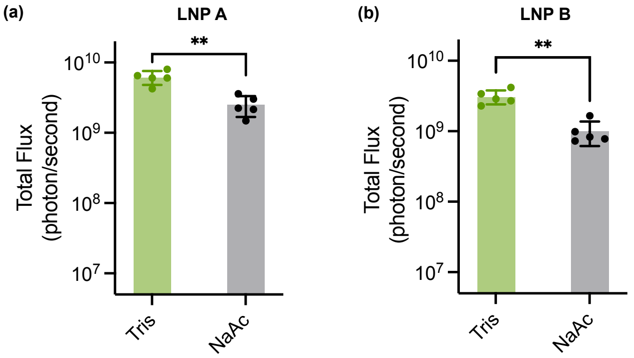


**Supplement Fig. 3 Effect of buffer on *in vivo* protein expression of solid-state FLuc mRNA-LNP formulations.** FLuc mRNA-LNPs formulated with the optimized excipient combinations were prepared in either Tris buffer (pH 7.5) or sodium acetate (NaAc, pH 5.5) for (a) LNP A and (b) LNP B lipid compositions.

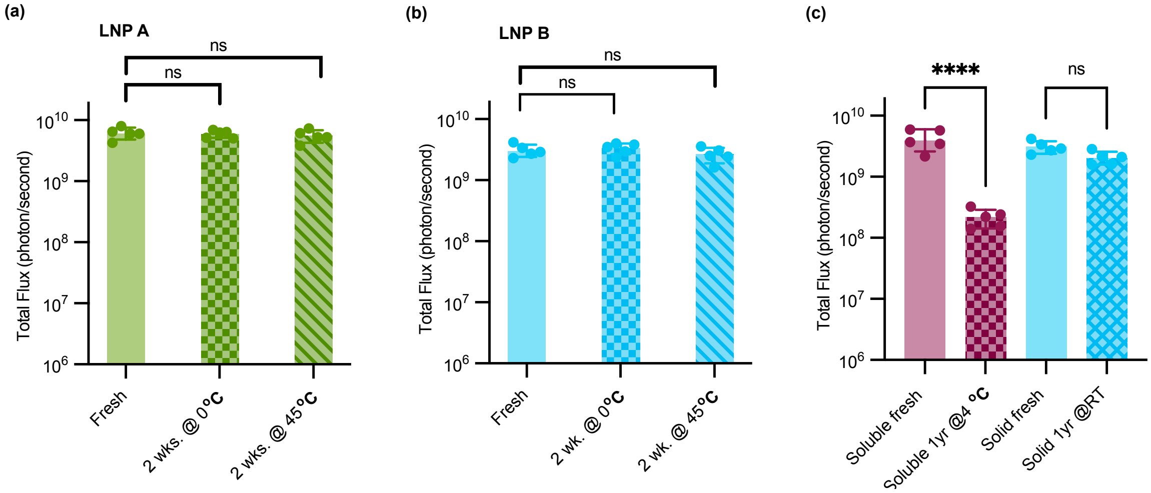


**Supplement Fig. 4 *In vivo* validation of solid-state FLuc mRNA-LNPs stability across storage temperatures.** Quantification of luminescence intensity at the injection site in mice at 6 h following intramuscular administration of reconstituted fresh or stored solid-state FLuc mRNA-LNPs (0.3 µg mRNA per mouse) for **(a)** LNP A and **(b)** LNP B. **(c)** Comparison of *in vivo* luminescence between soluble and solid-state FLuc mRNA-LNPs (0.3 µg mRNA per mouse, LNP B) after 1 year of storage. n = 5 biologically independent mice per group, one-way ANOVA with Tukey's test.

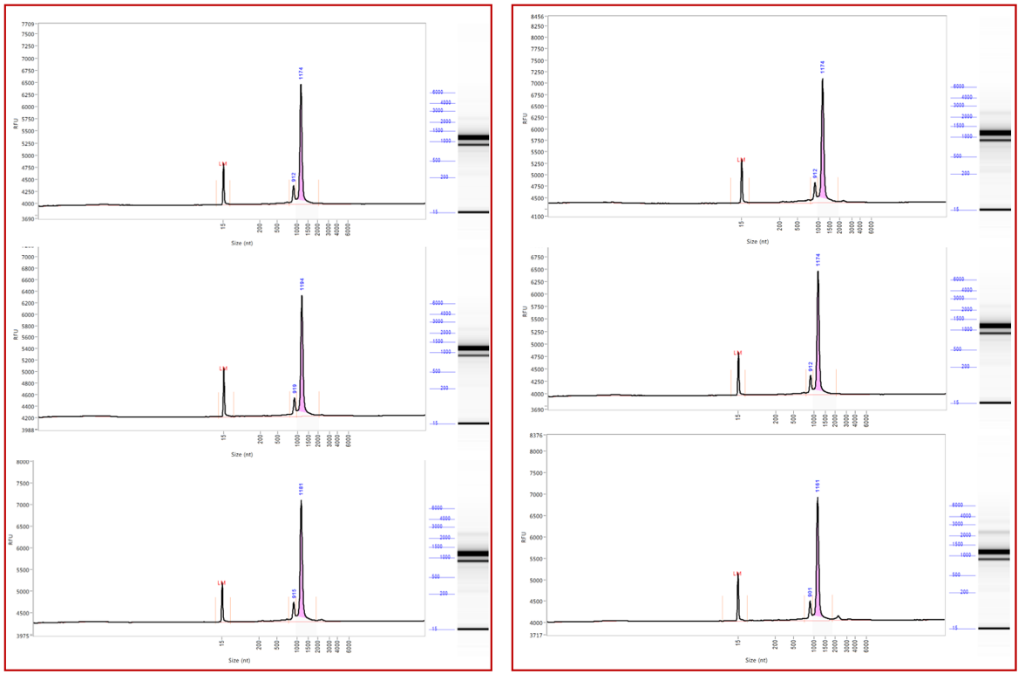

**Supplementary Fig. 5 FEMTO Pulse traces of soluble mRNA-LNPs and reconstituted optimized solid-state mRNA-LNPs.** Gel electrophoresis was performed on mRNA extracted from soluble LNPs (left panels) and reconstituted optimized solid-state LNPs (right panels).

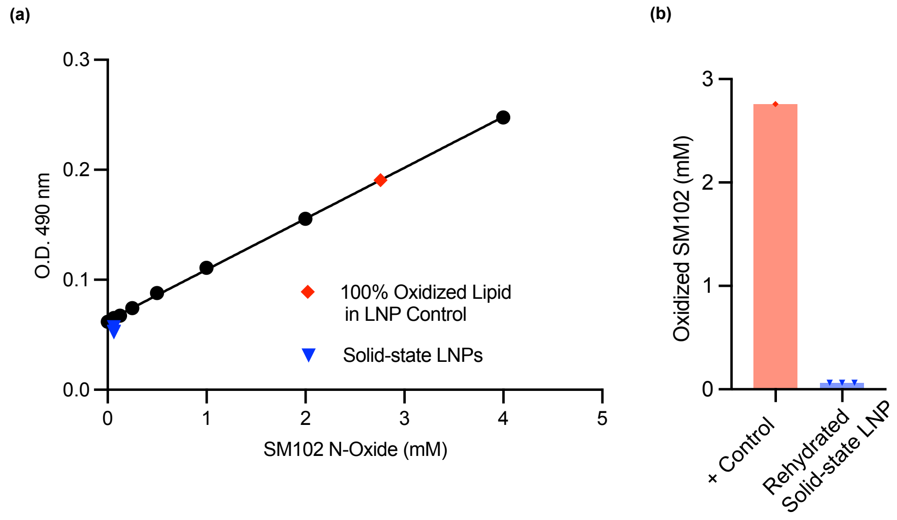


**Supplementary Fig. 6 Quantification of oxidized lipid in solid-state mRNA-LNPs.** (a) Standard curve generated using fully oxidized SM-102. (b) Concentration of oxidized SM-102 in 100% oxidized positive LNP control and reconstituted solid-state LNPs after vacuum drying.

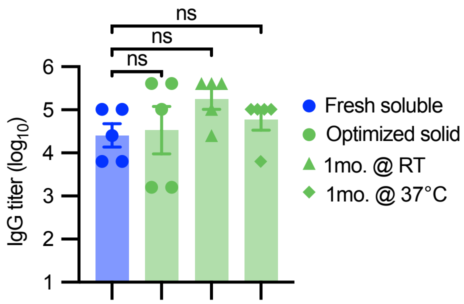


**Supplementary Fig. 7** Serum anti-RBD IgG titers measured at week 5 in mice administered fresh soluble or reconstituted solid-state RBD mRNA-LNPs that were stored at room temperature or 37°C for one month (LNP A lipid composition; n = 5 biologically independent mice per group, one-way ANOVA with Tukey's test.)

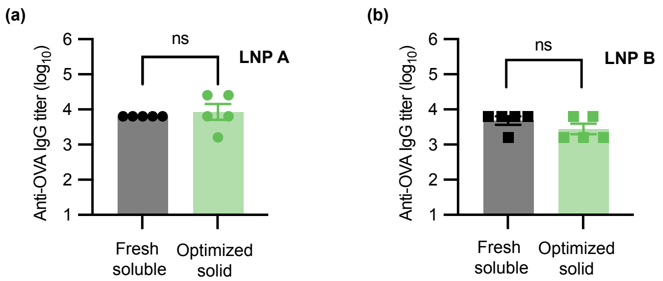


**Supplement Fig. 8** ***In vivo*** **validation of solid-state OVA mRNA-LNPs.** BL/6 mice were immunized with soluble or reconstituted solid-state OVA mRNA-LNPs via i.m. injections followed by a booster dose 2 weeks later. Plot shows serum anti-OVA IgG titers measured at week 3 for (a) LNP A and (b) LNP B lipid compositions (n = 5 biologically independent mice per group; 3 μg mRNA per mouse). Statistical significance was determined by two-tailed unpaired t test.

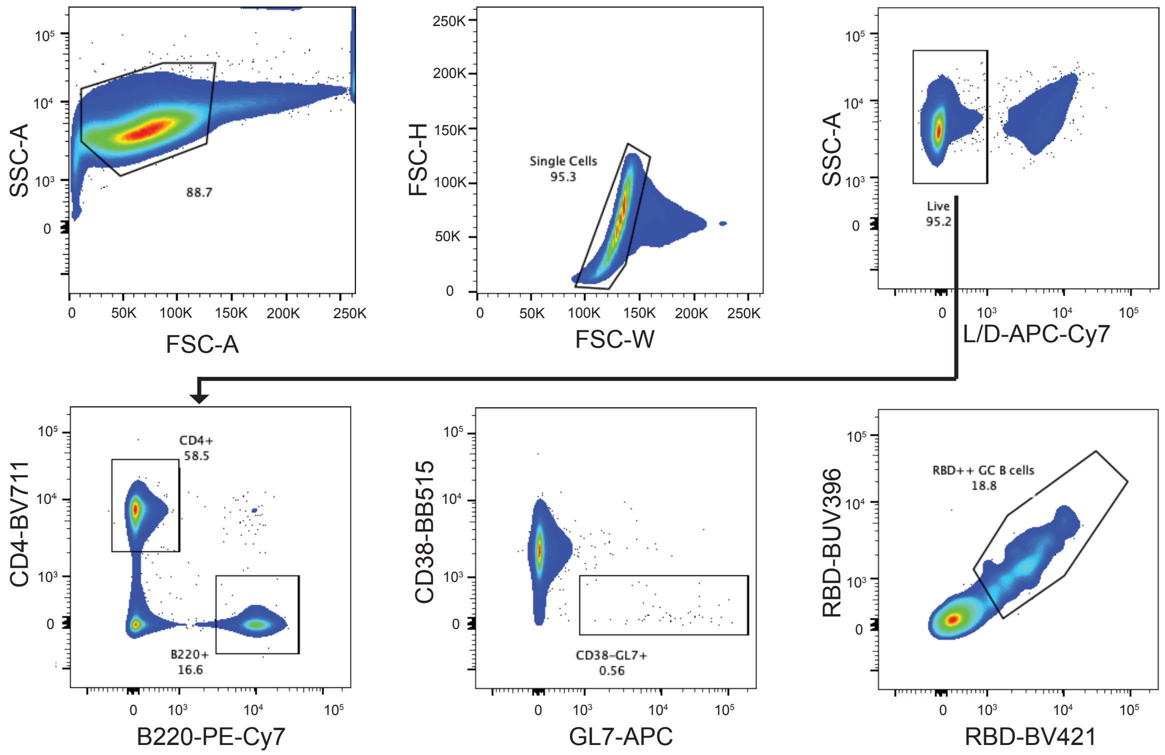


**Supplementary Fig. 9 Lymph node flow cytometry gating strategy.** Gating strategies for RBD-binding germinal center B cells derived from draining lymph nodes. Results are from one experiment. The gating strategy is reproduced in Fig. 4f.
